# Synchronization in collectively moving inanimate and living active matter

**DOI:** 10.1101/2022.10.06.511128

**Authors:** Michael Riedl, Jack Merrin, Michael Sixt, Björn Hof

**Affiliations:** Institute of Science and Technology Austria (IST Austria), Klosterneuburg, Austria

## Abstract

Regardless of whether one considers swarming insects, flocking birds, or bacterial colonies, collective motion arises from the coordination of individuals and entails the adjustment of their respective velocities. In particular, in close confinement, such as those encountered by dense cell populations during development or regeneration, collective migration can only arise coordinately. Yet, how individuals unify their velocities is often not understood. Focusing on a finite number of cells in circular confinements, we identify waves of polymerizing actin that function as a pacemaker governing the speed of individual cells. We show that the onset of collective motion coincides with the synchronization of the wave nucleation frequencies across the population. Employing a simpler and more readily accessible mechanical model system of active spheres, we identify the essential requirements to reach the corresponding collective state, i.e. the synchronization of the individuals’ internal oscillators. The mechanical ‘toy’ experiment illustrates that the global synchronous state is achieved by nearest neighbor coupling. We suggest by analogy that local coupling and the synchronization of actin waves are essential for emergent, self-organized motion of cell collectives.

## INTRODUCTION

Collective motion emerges from the self-organization of the individuals and occurs in living as well as inanimate active matter and across a wide range of length scales. Observations reach from macroscopic flocks of birds, or schools of fish, over the coordinate motion of swarms of robots^1,2^, down to the collective motion of cells^3^ and synthetic self-propelled particles.^4–8^ The most prominent model capable of reproducing e.g. flocking reduces the phenomenon’s complexity to a balance between the alignment strength between individuals and the noise disturbing the system.^9^ As in this and similar minimal models, the speed of individuals is assumed to be a constant. The effects of velocity fluctuations, which can destabilize the ordered state^10,11^, remain poorly understood. Especially in densely packed environments, like monolayers of migrating cells, feedback mechanisms between individuals are essential to ensure the coordination of the individuals’ velocities, a key requirement for the motion of the collective.

## RESULTS

We tracked the motion of cultured endothelial cells adhering on two-dimensional adhesive patterns shaped as disks or rings. When cells were plated at low density and had no contact with their neighbors, they moved without any preferential direction.^12^ When grown to confluency, cells transitioned from an initially chaotic motion to a coherently-moving state, where the collective adapted a carousel-like motion with uniform locomotion speed of the individual cells.^12–14^ **(Fig. 1a)**. Polymerization dynamics of the actin cytoskeleton provide the driving mechanical force for cell locomotion. We thus visualized actin dynamics by genetically introducing a fluorescent actin reporter in endothelial cells.^15^ Time-lapse fluorescent imaging revealed waves of high actin density traveling through cells **(Supp. Movie 1)**, a phenomenon previously described in other cell types and a consequence of the excitable dynamics of the actin polymerization machinery. ^16–19^ The formation of waves was evident in single cells as well as in collectives **(Supp. Movie 2)**. We initially confined single cells within a cell-sized circular adhesive pattern to quantify wave dynamics. From the resulting kymographs, we extracted individual waves propagating with an average speed of 9.4±2.7 μm min^-1^ **(Supp. Fig. 1a-f)**. To see the interplay between the internal actin waves and the speed of the cell, we restricted cells to one-dimensional movement on a line-like adhesive pattern. The nucleation frequency of newly arising waves within moving single cells varied from 2.2 to 5.5 h^-1^ and correlated positively with the speed of the migrating single cell **(Fig. 1d-e, Supp. Fig. 1g-h)**. That correlation was consistent with observations in other cell types where the nucleation frequency of actin polymerization waves correlated with the velocity of the leading edge. ^16,20–23^

**Fig. 1.**
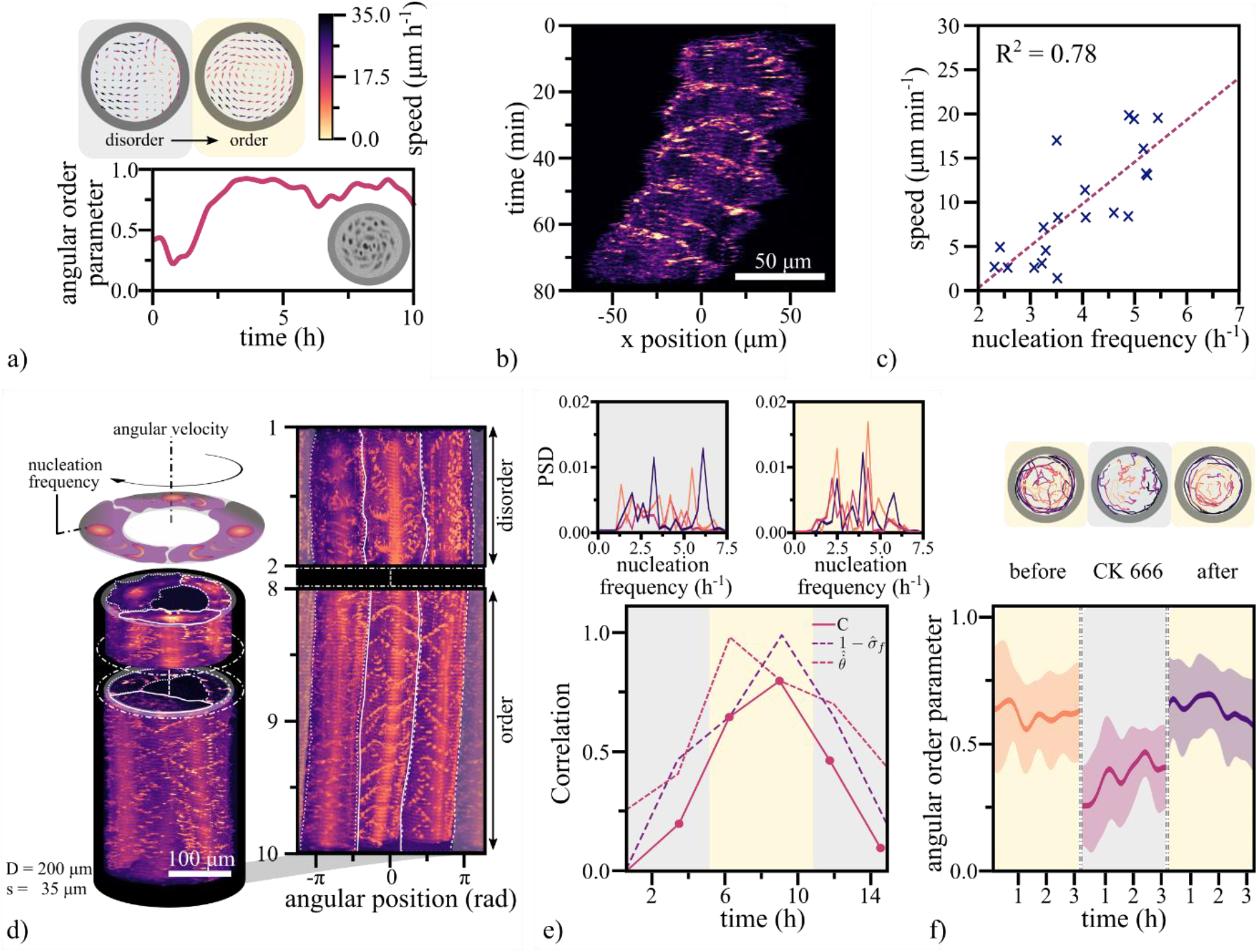
Transition to order in confined endothelial cells. **a**, Top: The state of disorder (grey shaded region) and order (beige shaded region) is illustrated by the corresponding velocity field of the confluent endothelial cell layer on a circular pattern (D = 200 μm). Bottom: The angular order parameter quantifies the transition from the initially disordered to the ordered motion. **b**, The space-time representation of a single cell moving on an adhesive line pattern (width = 50 μm. **c**, By plotting the mean migration speed of a single cell against the nucleation frequency of actin waves, we find a positive correlation (n = 20). **d**, Top: A schematic representation of the collective migration of endothelial cells on a ring-shaped adhesive pattern. Below: The space-time representation of a representative subset of the acquired time-series of a configuration of 3 cells on a ring (diameter = 300 μm, width = 35 μm), before and after the onset of collective rotation. Adjacent: In the corresponding polar transform, the frequency locked state becomes evident by the uniform vertical spacing between the periodically reappearing polymerizing actin waves with respect to the individual cells and across the collective. The white lines indicate the boundary between individual cells. **e**, Bottom: The correlation parameter *C* characterizes periods of small standard deviations in frequency and high mean velocities across the population. During phases of ordered, collective migration (beige shaded region), the correlation parameter C approaches values close to unity and decreases towards zero while the system is in disorder (gray shaded regions). Top: The corresponding power spectrum density depicts the dominant frequencies present. The main nucleation frequency of polymerizing actin waves is around 4.8 h^-1^, and the first harmonics are present at around 2.4 h^-1^. **f**, Top: Representative cell trajectories are shown during individual stages of a CK666 treatment experiment. During the presence of the ARP2/3 inhibiting drug, cells are actively migrating, but the global order is lost. After removing the drug, the carousel-like motion recovers. The transitions between states can be quantitatively identified by the drop and consecutive rise in the angular order parameter (n=10).

On the adhesive ring pattern that contained three cells, the individual nucleation frequencies were uncorrelated during periods where cells were not rotating collectively **(Fig. 1d-e, Supp. Movie 3)**. With the onset of collective migration and for as long as it persisted, the nucleation frequencies adjusted to a common frequency around 4.8 h^-1^ **(Fig. 1e)**. In non-rotating patterns, the nucleation frequency of polymerizing actin waves remained non-uniform across the collective **(Supp. Fig. 2)** By introducing the correlation parameter 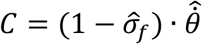, we quantified the correlation between the maximum in the normalized mean angular velocity 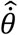 and the minimum in the normalized standard deviation of the frequencies 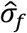 of the collective. We normalized both quantities by their respective maxima. We found that C reached its maximum during phases of ordered collective migration and decreased when the collective motion was arrested. Thus, the onset of collective motion coincided with the synchronization of nucleation frequencies **(Fig. 1e)**.

To probe whether collective locomotion and actin waves are causally related, we inhibited the chemical reaction generating the waves of polymerizing actin. Traveling actin waves are known to be driven by the self-activating WAVE protein complex.^19^ WAVE drives nucleation of actin polymerization via the Arp2/3 complex, and polymerized actin inactivates the WAVE complex, thus creating an excitable system.^17–19^ We pharmacologically perturbed Arp2/3 mediated actin nucleation with the drug CK666. We found that at a 100 μM concentration of the Arp2/3 inhibitor eliminated actin waves while single cell locomotion was maintained, albeit at lower velocities.^16^ The carousel-like rotations on adhesive surfaces were abrogated upon treatment with CK666, and cells migrated in an uncoordinated manner instead. Upon washout of the drug, the effect was reversed, and the collective rotation was restored **(Fig. 1f, Supp. Movie 5)**.^13^

Overall, our data suggest that collectively moving cells are coupled chemical oscillators driven by the excitable dynamics of the actin polymerization within each cell.^16–19^ We propose that intercellular coupling through nearest neighbor interaction gives rise to frequency locking, which in turn allows cells to adopt a uniform speed, enabling collective motion.

The observed collective migration that emerges from the synchronization of the individuals’ oscillators is reminiscent of recent theoretical models reproducing swarm formation.^24,25^ One such model introduces agents, known as swarmalators^24^, whose velocity is expressed as a function of the phase of an internally oscillating process. By further introducing coupling between the oscillators, they potentially synchronize and the collective transitions from an uncoordinated motion to a swarm-like state. To experimentally test if nearest neighbor coupling suffices to establish synchronization as an underlying mechanism to unify velocities and eventually lead to collective motion, we introduced a model system of mechanical ‘toys’ with comprehensive degrees of freedom. We employed ‘weaselballs’ (D.Y. Toy), autonomously moving spherical shells driven by an internal, unbalanced motor freely rotating around an axis connecting the poles of the shell. The motor applies a constant torque and while the shell is fixed causes the continuous revolution of the internal mass **(Supp. Fig. 3a)**. We extracted the frequency of the revolution by tracking the position of the motor over time and found that it varies between individual balls **(Fig. 2a, Supp. Fig. 3b,c)**. When the motorized balls are released on a sufficiently rough substrate, they roll with oscillating speed along unstable trajectories **(Fig. 2b)**. After coming into contact with the boundary of the circular confinement, a single ball traces it, a behavior common for active particles **(Supp. Fig. 2b)**.^1,26^ Along the resulting circular trajectory of the motorized ball the speed oscillates **(Supp. Fig. 3d)**. The frequency of the oscillating speed is reduced with respect to the frequency of the corresponding freely rotating motor **(Supp. Fig. 3e-f).**

**Fig. 2.**
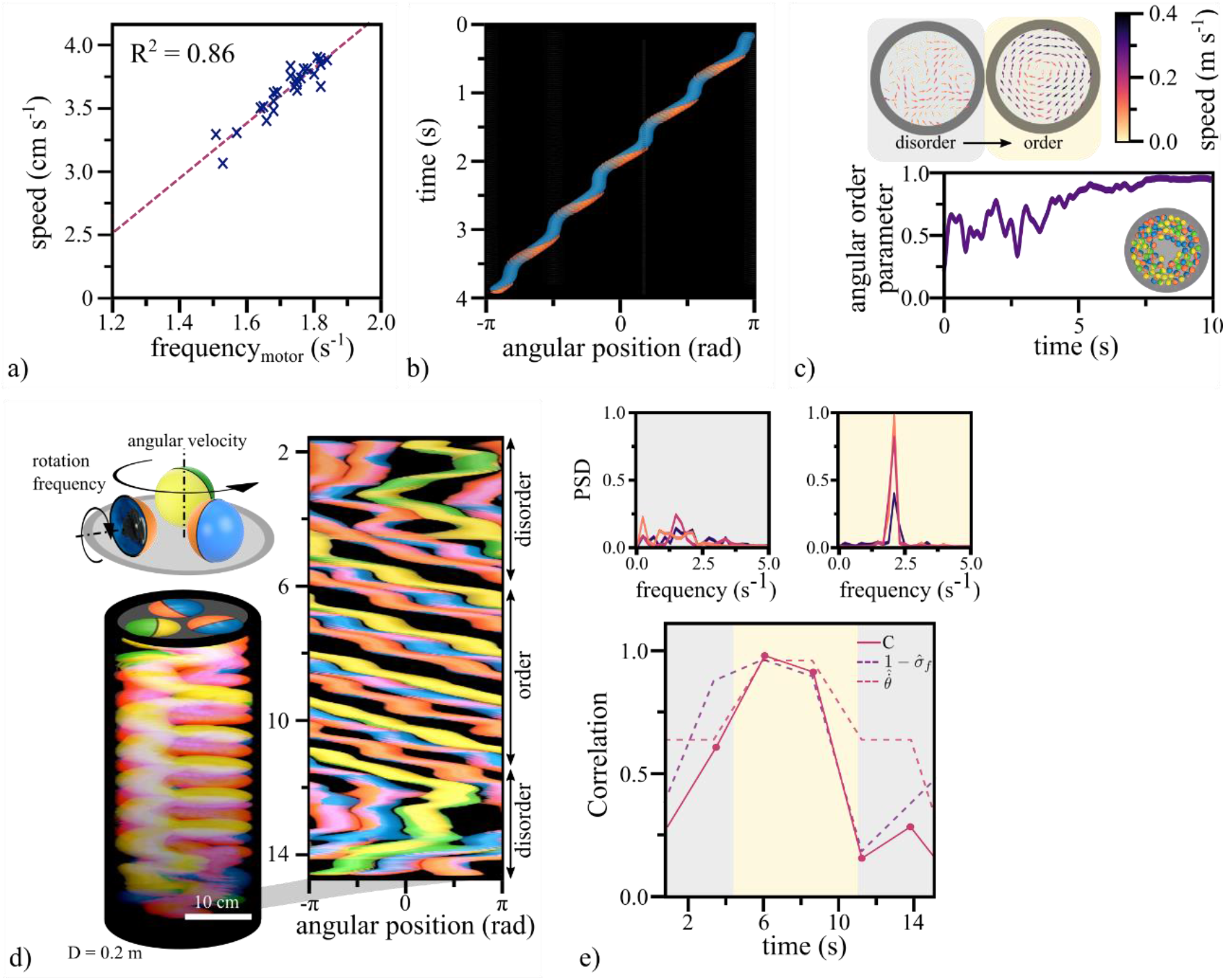
Transition to order in confined motorized balls. **a**, The mean speed of a single motorized ball correlates positively with the frequency of the internal motor, which was measured in an opened ball while the shell was fixed (n = 25). **b**, The staircase-like trajectory in the space-time representation visualizes the oscillatory motion of a single ball in a circular confinement. **c**, Top: A representative velocity field of the motion of a population of motorized balls (N = 73) in a circular confinement (D = 0.9 m) in the state of disorder (grey shaded region) and order (beige shaded region). Below: The angular order parameter quantitatively shows the spontaneous transition from the initial disordered to the ordered motion quantified. **d**, Top: The schematic depiction of a system containing three motorized balls. Below: The space-time representation of a subset of the acquired images. Adjacent: In the corresponding polar transformation, the synchronized state is recognizable by the periodically repeating, parallel paths of the ball trajectories. The slope indicates the uniform speed within the collective. **e**, Top: Plotting the power spectrum density in both states illustrates the transition between disorder and order. The initially broad distribution of frequencies converges to defined peaks around 1.9 s^-1^. Below: The correlation parameter C reaches values close to unity in the synchronized state, where the mean velocity is a maximum, and the standard deviation of the frequency maxima reaches a minimum.

When we placed multiple balls in circular confinements, their dynamics transitioned from a disordered motion to a smooth and ordered collective rotation pattern **(Fig. 2c, Supp. Movie 4)**. Once established, the rotational movement for large numbers of balls remained stable. In contrast, a confinement containing three randomly selected balls alternated between ordered and disordered states, allowing us to analyze the transition between these states **(Fig 2d, Supp. Movie 4)**. When we computed the angular velocity of each ball with respect to the center of the circular confinement, we found that the velocities oscillated out of sync during periods of disorder and in sync during the ordered state. Additionally, during the period of the ordered state, the mean angular velocity of the population 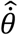 showed a maximum. This maximum overlapped with a minimum in the standard deviation of the instantaneous frequencies 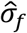 of the oscillators. We quantified this correlation with the previously introduced correlation parameter *C* and found that it reached values close to one during periods of synchronized motion and strongly decreased when the system was in disorder. **(Fig. 2e)** This shows that the onset of collective motion coincides with the locking of motor frequencies.

Synchronization requires coupling between the individual oscillators.^27,28^ In our system, coupling arose from the contact between balls. To investigate the extent of the coupling, we extracted the motor frequencies from trajectories of single balls individually moving in a confinement. We found a distribution of frequencies, between 1.3 and 1.9 s^-1^ **(Supp. Fig. 3e)**. The same population of balls, when put together in a circular or annular confinement, transitioned to a coherently rotating state. In this state, the initially wide frequency distribution collapsed to a single, common frequency **(Supp. Fig. 4)**. This indicated that the coupling mechanism did synchronize not only the phases of oscillating motion of rolling balls but also the frequencies of the motors. To establish a clear picture of this process, we produced a set of balls with transparent half-shells and tracked the motion of the motor independent of the outer shell **(Fig. 3a,b)**. From the oscillations of each motor, we extracted the frequency spectra **(Fig. 3c)** which confirmed that the motor frequencies locked by converging to a common frequency **(Fig. 3d)**. The observed synchronization originated at the level of the motors and extended to the movement of the balls.

**Fig. 3.**
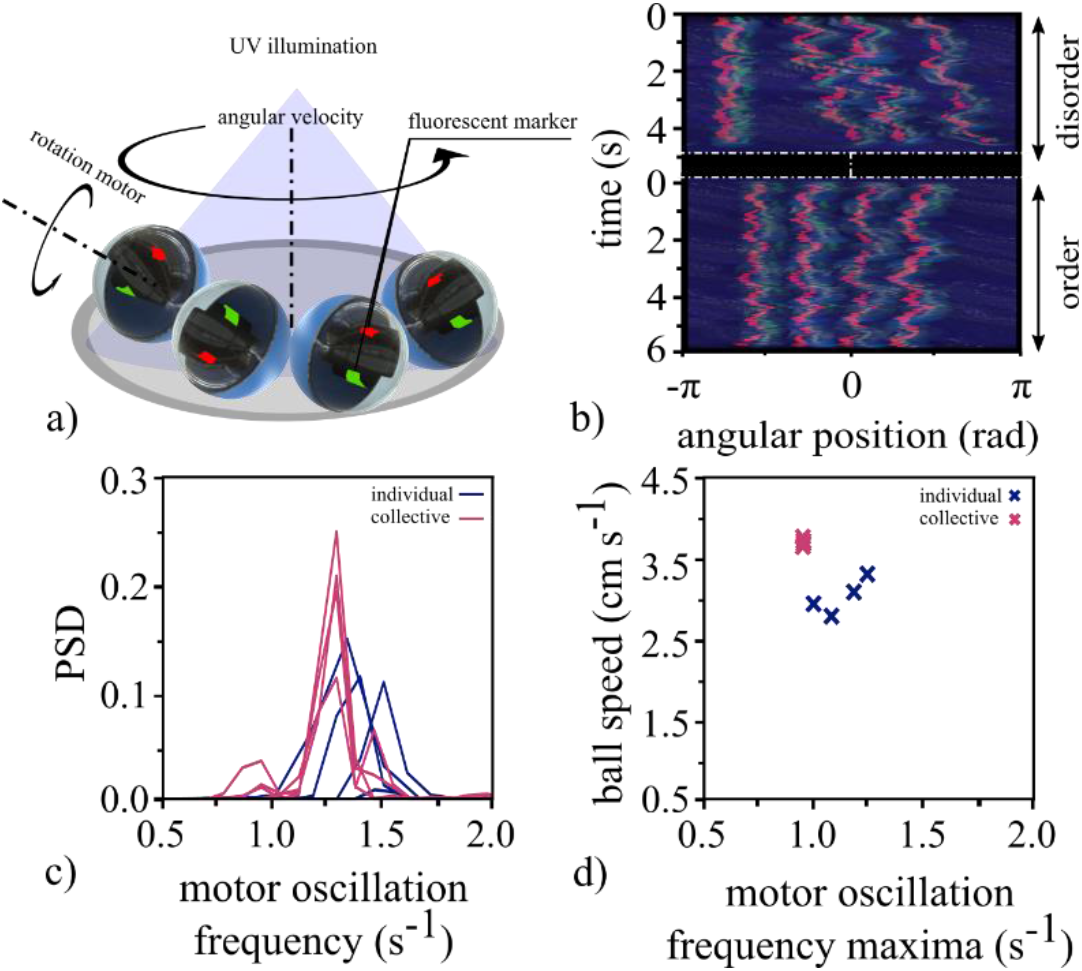
Experimental setup and synchronization of the motors in confined balls with transparent half-spheres. **a**, The schematic representation of the experimental setup using 4 balls with transparent half-spheres in a circular confinement (D = 0.4 m). We attached fluorescent tape to the internal motors and illuminated the setup with UV light to visualize the motion of the internal motor. **b**. We show the space-time representation of a disordered and ordered period in the comoving reference frame with respect to the leading ball. The red oscillations of the fluorescent tape correspond to the oscillation of the internal motors. During periods of order, the oscillations synchronize their frequencies and phases. **c**, The corresponding power spectrum density shows this change in the frequency distribution of the internal motors’ oscillation. **d**, The oscillations of the internal motors become independent of the balls’ motion by stabilizing it with respect to the velocity corresponding to the individual ball’s center of mass. Plotting the frequency maxima of these oscillations shows that the frequencies lock during synchronized movement.

We next investigated mechanisms to destabilize the ordered state or to prevent it from arising. We probed this by continuously increasing the number of balls within a confinement (D = 0.9 m) moving in an ordered state. After reaching a critical number of agents (N = 82) in the confinement, the ordered motion of the now densely packed population broke down and transitioned into the disordered state. The transition to disorder corresponded to a sudden drop in the order parameter **(Fig. 4a, Supp. Movie 5)**. When the population of balls was preselected to have matching natural frequencies to avoid variability within the collective, the ordered state still emerged in fully packed confinements **(Supp. Movie 5)**. This indicated that the stability of the ordered motion depended on the distribution of frequencies rather than the density of the population. As shown previously^29^, collective motion is sensitive to uncooperative individuals or variations within the population. In flocks, already a low fraction of non-aligning agents can disrupt the collective movement.^29,30^ To see if this holds true for our systems, we first probed our system of motorized balls by exchanging active with inactivated balls by switching their motors off. The substitution of about 4% of the population with uncooperative balls led to the breakdown of the collective behavior, in quantitative agreement with values found in numerical simulations on flocking.^29^ **(Fig. 4b, Supp. Movie 5)**

**Fig. 4.**
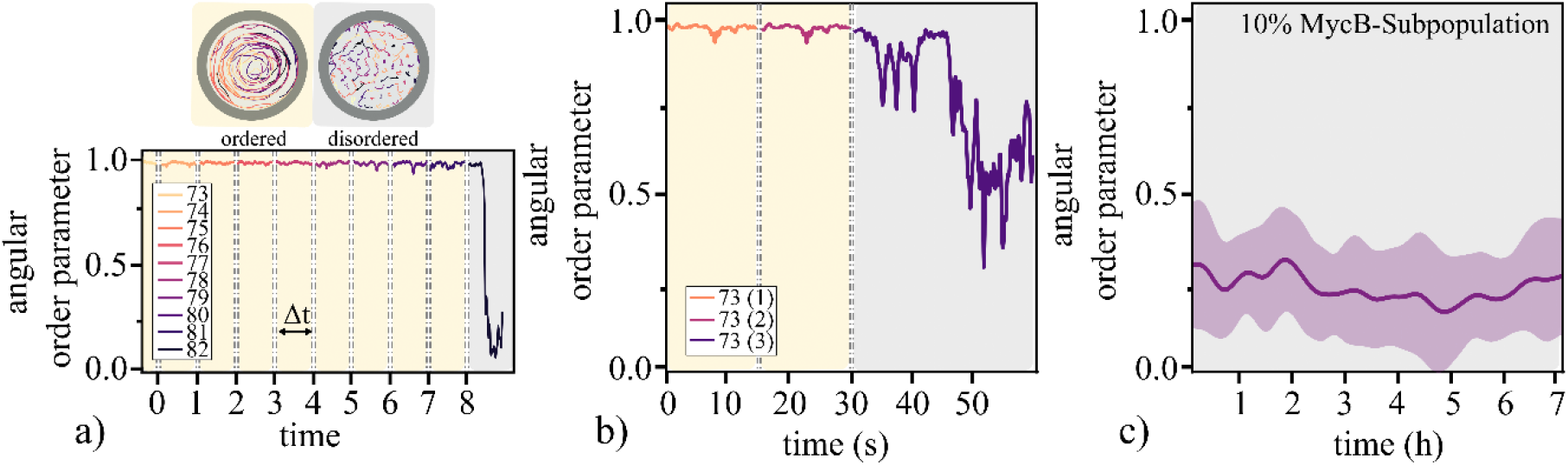
Destabilizing the order in both systems. **a,** We approached the critical number of agents at n = 82 by incrementally increasing the number of balls in a circular confinement (D = 0.9 m). Top: Representative trajectories are displayed for both states over an equal time interval of 2 s. They show the transition from a collectively moving state to a Brownian-like motion. Bottom: The steady states’ angular order parameter fluctuates close to unity for a representative time interval of Δt = 25s. The transition to disorder is evident by the sudden decrease in the angular order parameter when reaching a critical number of agents within the confinement. **b**, Consecutively rendering 4 % of the population of balls inactive leads to the breakdown of the order (3 inactive balls within a collective of 73 balls). **c**, Partially inactivating actin polymerization in a subpopulation of cells prevents order from arising (n=12).

To see whether our findings on destabilization in our system of motorized balls are translatable to our system of migrating cells, we aimed for the conceptually analogous experiments. In agreement with our observations in the system of motorized balls, previous findings show that the rotating motion of migrating cells in confinement is arrested after reaching a critical density threshold.^14^ To perform the experiment analogous to introducing uncooperative individuals, we chose to treat endothelial cells with the actin polymerization inhibitor MycB at doses that inhibit actin wave formation but not adhesion and spreading on the substrate. Contrary to most other pharmacological agents, MycB binds covalently to actin and therefore does not wash out upon dilution or leak into other cells. We pre-incubated a subpopulation of endothelial cells with MycB and simultaneously labeled it with a fluorescent dye so that it could be identified and tracked when mixed with untreated cells. MycB renders the subpopulation of treated cells into passive individuals that were dragged around by their active neighbors. The presence of these uncooperative individuals prevented the ordered state from arising, in agreement with our observations in the mechanical experiment as well as models simulating flocking^29^ **(Fig. 4c, Supp. Movie 5)**.

## DISCUSSION

In nature, propulsion mechanisms are often based on repetitive processes like the flapping of wings or fins or the rotation of cilia. For the collective motion to arise and persist, feedback mechanisms are required to enable these autonomous oscillators to adjust^31,32^ and converge to a uniform speed^30^. As proposed in the original theoretical work on swarmalators^24^ and demonstrated in the systems presented here, weak coupling between the underlying oscillators is a sufficient and conceptually simple way to establish the necessary feedback. Eventually, the emerging synchrony results in the uniformity in speed required for collective motion. Usually, this process relies on neighbor interactions, transferring information across the collective.^33^ The speed as well as the robustness of this transfer in turn limits the size of the coherent collective motion.^34^ This limitation potentially explains the similarity in length scales of collectively rotating cells observed here and those found in spontaneously emerging transient vortices appearing in larger confinements or unconfined monolayer of otherwise chaotically migrating cells.^12^ As we show, the propagation process can be compromised by variations between the oscillators or the presence of uncooperative individuals destabilizing the arising order. This sensitivity of collective motion limits the variability between entities and restricts it to contributing individuals. While the overall concept is not restricted to the unicellular or mechanical agents studied here^35^, it remains to be seen whether the collective motion of more complex organisms relies on similar principles.

## Methods

### Cell culture

#### Lentivirus production

The lentivirus production was performed as previously described^20^. In brief, LX-293 HEK cells (Clontech) were co-transfected with LentiCRISPRv1 or LentiCRISPRv2, packaging-psPAX2 (Addgene no. 12260) and pCMV-VSV-G envelope plasmids using Lipofectamine 2000 (Thermo Fisher Scientific) as recommended by the manufacturer, and cells were resuspended in R10 medium 1 day before transfection. The supernatant was collected after 72 h and stored at −80 °C.

#### Cell Culture

Human Aortic Endothelial Cells (HAoEC), obtained from PromoCell GmbH, were cultured in growth medium (Endothelial Cell Growth Medium MV, PromoCell). Cells were grown and maintained in a humidified incubator at 37 °C and 5% CO2 and passaged after reaching 70% confluency. During long-term imaging, we added 1% penicillin–streptomycin antibiotic (Invitrogen) to the growth medium.

#### Cell infection and selection

For the infection of HAoEC with the Lifeact-GFP reporter construct^15^, we use a 70 % confluent monolayer in a T-75 flask of a passage number not higher than 2-3. After aspiration of the initial growth medium, we added 10 ml growth medium supplemented with 0.5 ml Lentivirus and incubated overnight. Subsequently, we removed the growth medium and washed the cell layer 3 x with PBS before adding fresh growth medium. Afterward, we proceeded by starting the selection process by supplementing 3 μl Blasticidin (Gibco, Thermo Fisher Scientific) from a stock solution (10 mg/ml) to 10 ml cell culture medium. The selection is completed after 4-6 days or until a sufficiently low number of non-fluorescent cells remain.

#### Photomasks

The patterns were designed with Coreldraw X8 (Corel Corporation) exported to DXF, then converted to GERBER format with LinkCad. Chromium quartz photomasks (100 mm or 125 mm; PhotoData/JD Photo-Tools) were used.

#### Patterns

The adhesive patterns were generated using PLL-g-PEG (Surface Solutions, Switzerland) as previously described^21^ with a few changes. Initially, we rinsed the coverslips with ethanol and dried them with filtered compressed air. After activation by exposure to UV light for 15 min, we incubated the coverslips with 0.3 mg/ml PLL-g-PEG for 1h at 37°C, which created an inert, non-adhesive coating. Subsequently, UV irradiation through the chromium quartz photomask removes the non-adhesive coating locally, creating the desired pattern. Before seeding the cells, we washed the coverslips 1x with Ethanol and subsequently 2x with PBS and coated them with 0.2% gelatin.

#### Microscopy

Throughout the image acquisition process, the cells were within an incubation chamber under a controlled CO2 atmosphere. Timelapse fluorescent images were recorded using an inverted wide-field Nikon Eclipse Ti-2 microscope equipped with a DS-Qi2 monochrome camera and a Lumencor Spectra X light source (475 nm Lumencor). NIS Elements software (Nikon Instruments) was used for acquisition control for e.g., multi-positions imaging.

#### CK666 treatment

After an initial unperturbed period, we treated the cells by adding a 100 μM solution of the ARP 2/3 inhibitor CK666 in preconditioned growth medium to the sample. Subsequently, the effect was reverted by removing the medium and washing the cells 2x with preconditioned growth medium. Each consecutive phase of this treatment was followed by a stabilization period of 4h before image acquisition.

#### MycB treatment and TAMRA staining

We prepared the staining solution by diluting 1 ul of a 10 mM stock solution of TAMRA in 1 ml PBS. Subsequently, we washed the adherent HAoEC monolayer 1x with PBS and added the previously prepared staining solution. After an incubation period at room temperature of 10 min, we washed the cells 2x using preconditioned growth medium. In the next step, we diluted the MycB stock solution to 0.1 μM in growth medium and added it to an equal amount of growth medium in the flask with the MycB treated cells resulting in a final concentration of 0.05 uM. After a 30 min incubation period, the cells were gently washed 3x with preconditioned growth medium. After a recovery period of 30 minutes, cells were detached, mixed together with non-treated cells, and seeded on micropatterns.

### Weaselballs

#### Preparation

We removed the ‘weaselballs’ (D.Y. Toy) from their shipment boxes and separated them from the attached weasels. The balls consist of two half-spheres which can be screwed open to detach them, and after, one AA battery can be inserted. Before each experiment, the AA batteries had to be collectively exchanged to ensure similarity in the torque generation throughout the populations. Empty or nearly empty batteries can lead to artifacts in the rolling behavior of a ball.

#### Transparent half-spheres

We used silicone rubber (Wagnersil 22 NF) to create an inverse mold of the outside and inside of the desired half-sphere. Subsequently, we removed the half-sphere from the mold, mixed the transparent epoxy resin adhesive with the hardener (Let’s Resin), and filled the resulting gap. Previous to pouring, the two components had to be well mixed and free of air bubbles.

### Image analysis

FIJI imaging processing software (https://fiji.sc/) was used for image and video microscopy processing and analysis.

#### Trajectory Tracking

For extracting the trajectories of the balls or the cells from recorded movies, we used the ‘manual’ or ‘semi-automated tracking’ feature provided within the plugin TrackMate within the imaging process software FIJI.^36^ We post-processed the extracted trajectories using Python 3.7.

#### Phases and frequencies balls

From the extracted trajectories of the balls, we calculated the angular velocities with respect to the center of the confinement. From these oscillating velocities, we could extract the instantaneous phases and frequencies using the Hilbert transform method provided within the scipy library.

#### Frequencies Cells

For extracting the nucleation frequency from the kymographs generated with FIJI, we corrected the instantaneous position of each cell by the mean angular speed of the population. In the case of ring patterns, this step was precluded by a polar transformation of the initial images. Transforming the images in a cell-comoving reference frame results in a kymograph equal to a stationary pattern.

Subsequently, we use the Fast Fourier Transformation method within the scipy library to calculate the frequency spectrum at multiple positions within a cell over the time frame of interest. Each position within a cell results in a frequency spectrum, from which the global maximum of the dominant peak value across all calculated frequency spectrums serves as a selection criterion.

---

**Supp. Fig. 1.**
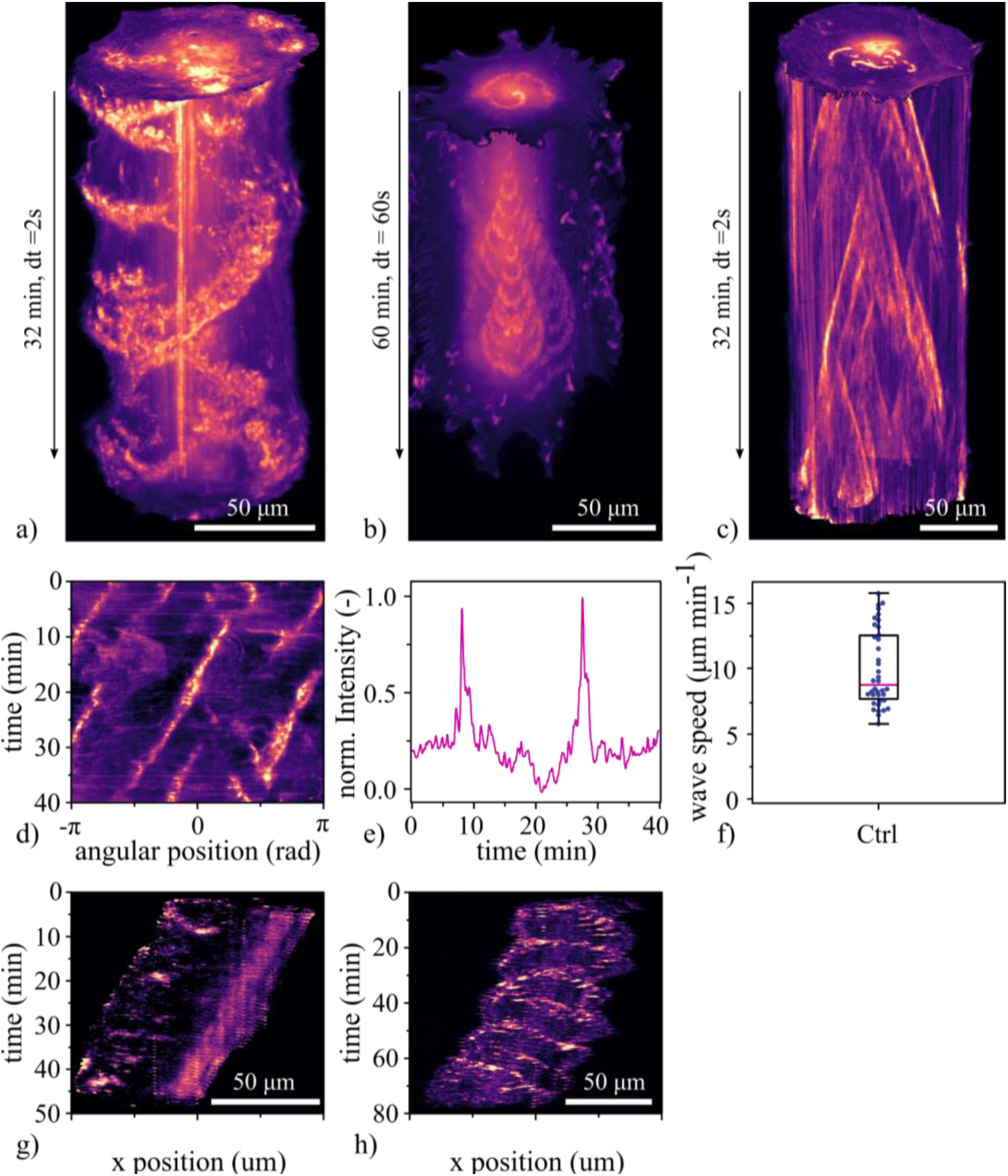
Polymerizing actin waves characterization in single cells on circular and line patterns. **a,b,c,** Space-time representations of different manifestations of the observed actin polymerization waves. While in **a** multiple persistent waves travel orderly through the cell, in **c**, small short-lived wavelets form and propagate through the cell. In **b**, we observe spiral wave formation reemerging from the same initial point. **d,** The polar transformed and sliced space-time representation of the wave dynamics corresponding to **a**. Here, up to 3 waves are present simultaneously and annihilate upon collision. **e,** Extracting the intensity along fixed angular position results in peaks corresponding to actin polymerization waves passing through. **f,** Actin polymerization wave propagation speed within stationary cells in circular confinement. **g, h,** Space-time representation of a single cell migrating on a line pattern (width = 50 μm). The polymerizing actin waves at higher **g**, and lower **h,** frequencies.

**Supp. Fig. 2.**
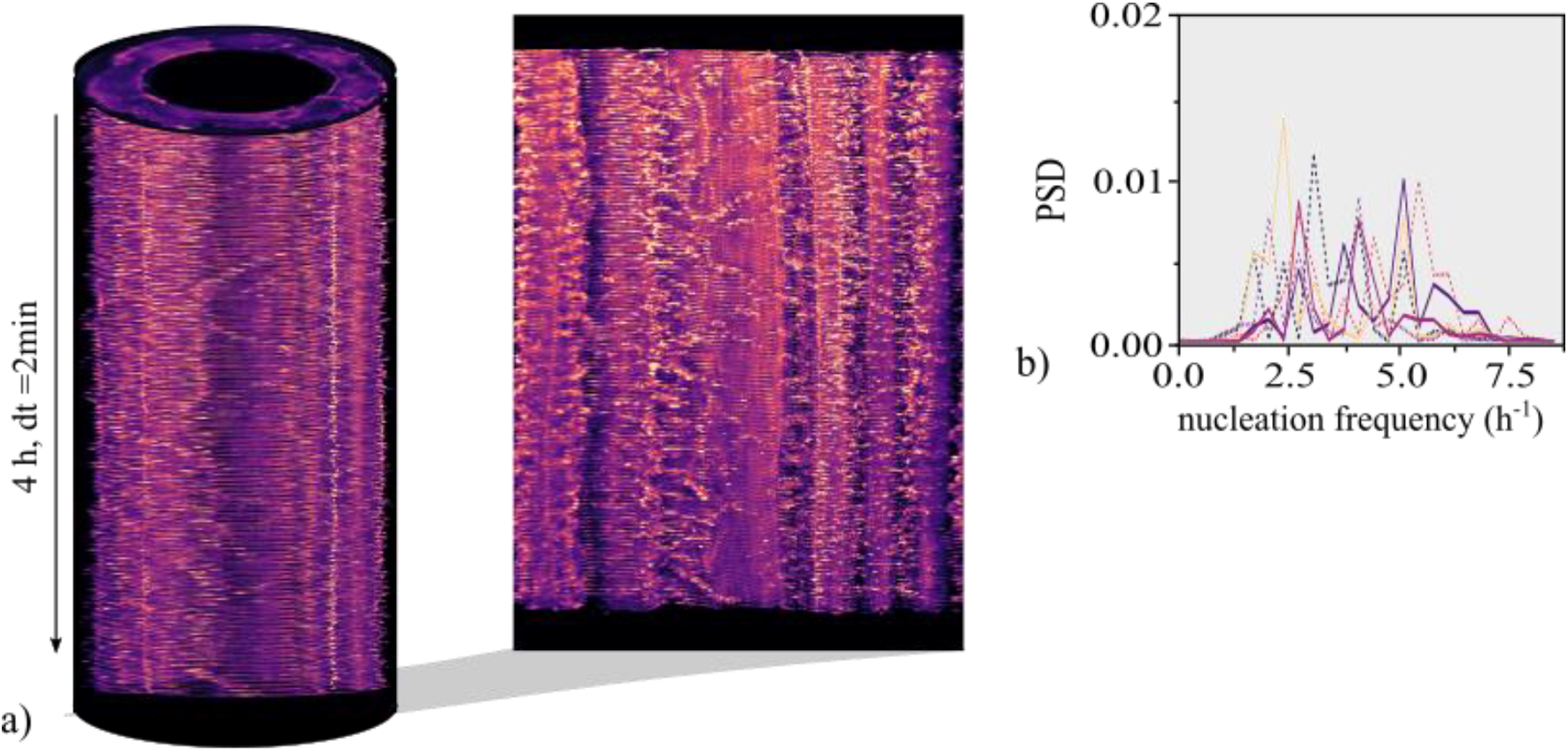
Absence of frequency locking and collective rotation. An example of 7 cells on an adhesive ring pattern (diameter = 300 *μ*m, width = 35 *μ*m) that do not collectively migrate. **a,** The space-time representation and the corresponding polar transformation omits collective rotation or convergence of nucleation frequencies of polymerizing actin waves. **b,** The power spectrum density plot omits a common dominant frequency across the collective, despite the presence of cell-specific frequency maxima.

**Supp. Fig. 3.**
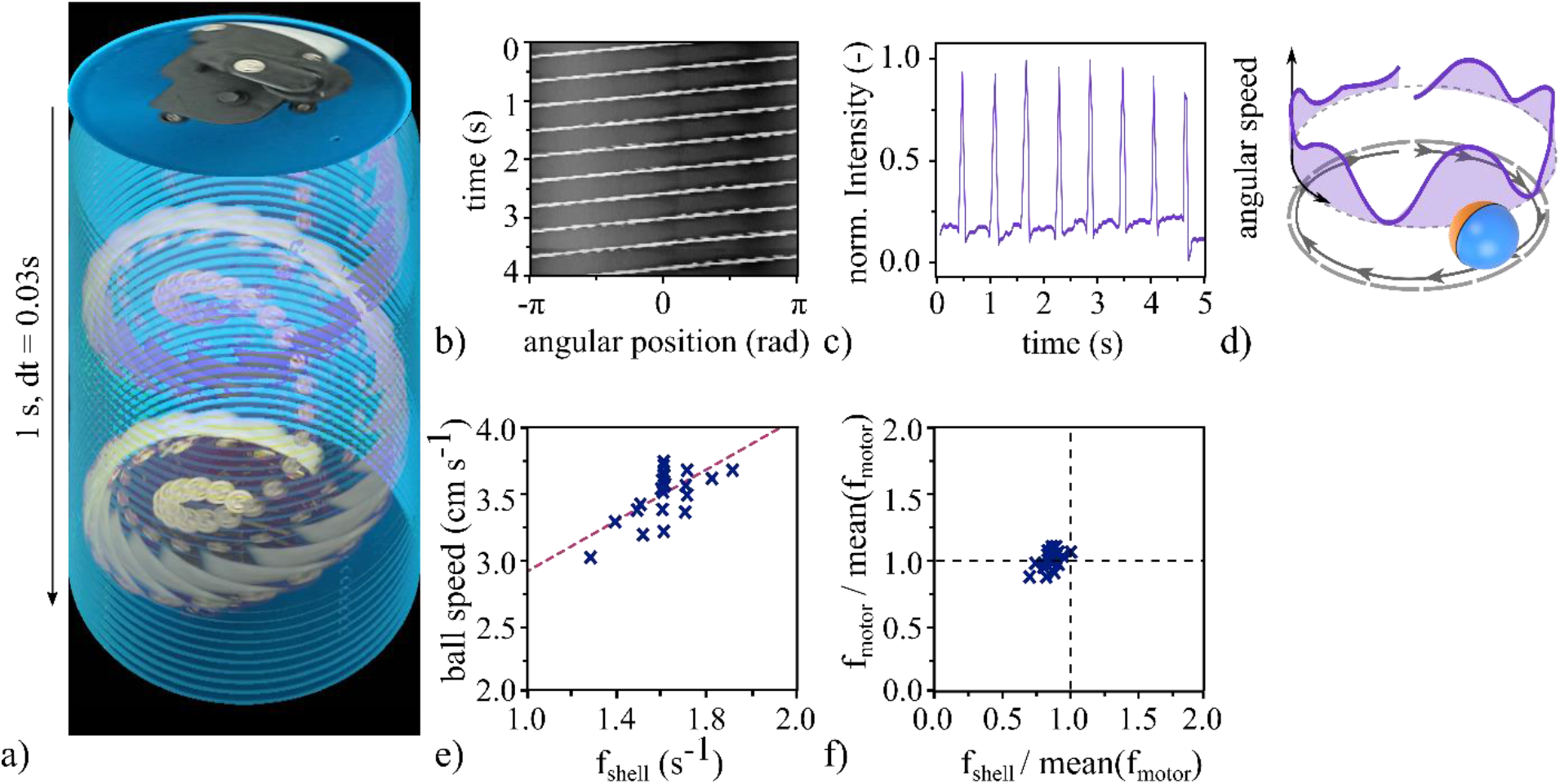
Single motorized ball characterization. **a,** The space-time representation of the freely rotating unbalanced motor within a fixated shell visualizes the rotating motion of the motor while the shell is being fixed. **b,** The position of the internal motor and weight periodically rotates in the fixated state around its axis. **c,** The extracted intensity peaks correspond to the revolutions. **d,** An example trajectory of a rotating ball shows the corresponding characteristic oscillations in speed along the boundary of the confinement (D = 0.2 m). **e,** The frequencies extracted from velocities of single balls rolling in a circular confinement show the non-uniform distribution (D = 0.5m, n =25). **f**, The rotation frequencies of the shells correspond to a negative shift of the motor frequencies. Both quantities were normalized with the mean motor frequency.

**Supp. Fig. 4.**
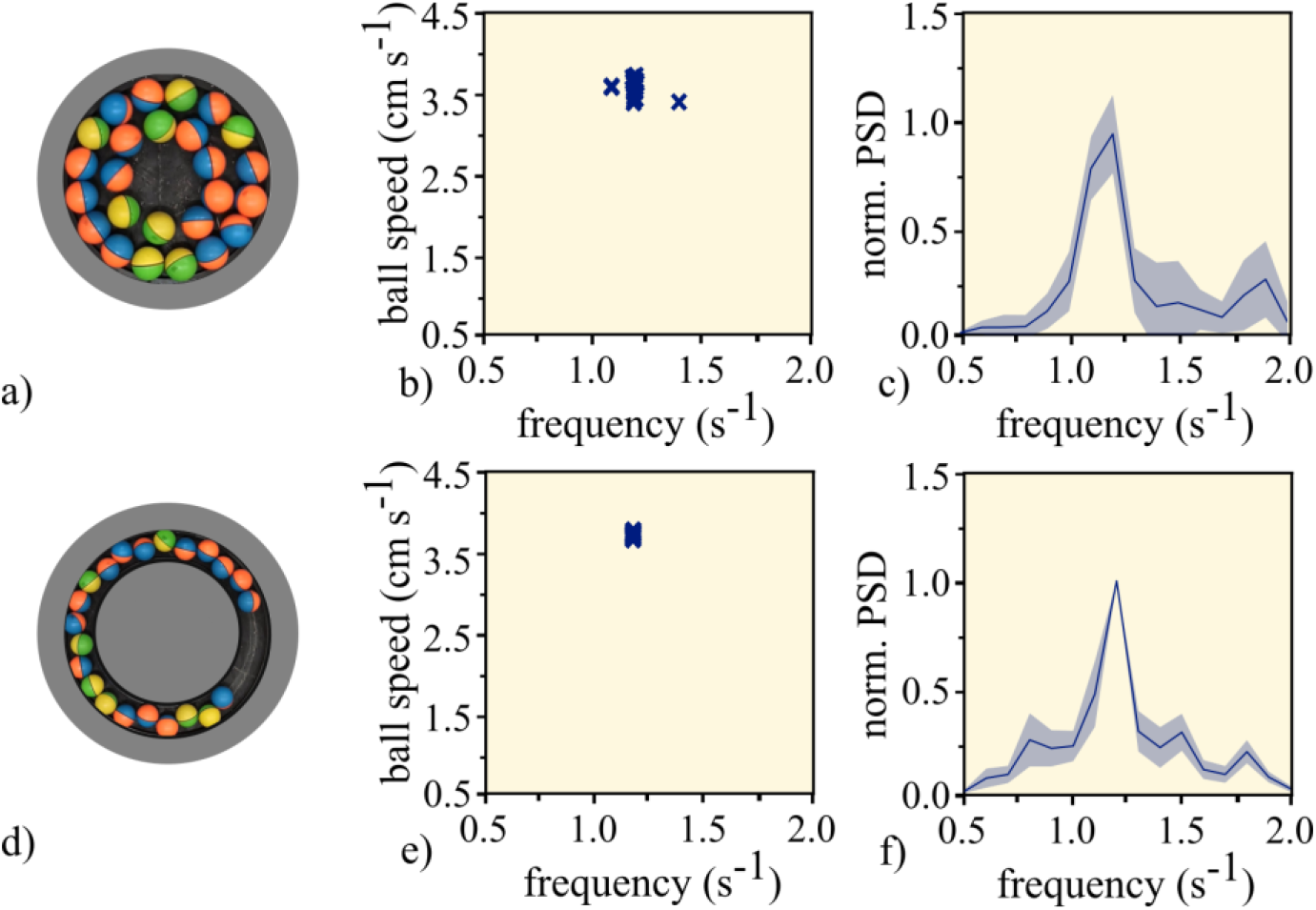
Rotation frequency characterization in collectives of motorized balls in circular and annular confinements. **a**,**d**, Snapshot of the ordered state in a **a**, circular confinement (D = 50 cm, n = 25) and **d**, ring confinement (D_outer_ = 75 cm, D_inner_ = 50 cm, n = 22). **b**,**e**, Frequency maxima in the ordered, collectively rotating state **b**, circular and **e**, ring confinement. **Fig. 2b**. shows the frequency of individual balls. During the ordered rotation, the individually spread frequency lock to a single collective frequency. **c,f**, The corresponding frequency spectrum density for the collectively rotating **c**, circle, and **f**, ring.

